## Supplementary Figure 1 for "Cyclic AMP signalling and glucose metabolism mediate pH taxis by African trypanosomes"

a

| Plate | left | right |
|---|---|---|
| 1 | Alanine | Proline |
| 2 | Threonine | N-acetyl-D-glucosamine |
| 3 | Fumarate (di-Sodium salt) | NaOH |
| 4 | FeCl3 | FeCl2 |
| 5 | LiOH | NaOH |
| 6 | Calcium acetate | HCl |
| 7 | Ammonium hydrogen sulfate | Ammonium hydrogen carbonate |
| 8 | Calcium hydroxide | Magnesium hydroxid carbonate |
| 9 | Iron(II) sulfate | Ammonium iron(II) sulfate |
| 10 | Ammonium acetate | HCl |
| 11 | KOH | NaOH |
| 12 | Cadmium acetate | HCl |
| 13 | Tri-potassium citrate | HCl |
| 14 | Magnesium acetate | HCl |
| 15 | Copper sulfate | NaOH |

b

| Attractant | pH | Repellent | pH | no visible effect | pH |
|---|---|---|---|---|---|
| NaOH | 14 | HCl | 1 | Calcium hydroxide | 12-13 |
| LiOH | 14 | FeCl3 | 2 | Ammonium acetate | 7 |
| KOH | 14 | FeCl2 | 2 | Copper sulfate | 3 |
| Ammonium hydrogen carbonate | 8-9 | Iron(II) sulfate | 2-3 | N-acetyl-D-glucosamine | 4-5 |
| Magnesium hydroxid carbonate | 9 | Ammonium iron(II) sulphate | 3 | | |
| Tri-potassium citrate | 7-8 | Ammonium hydrogen sulphate | 1 | | |
| Calcium acetate | 8 | Sodium acetate | 5-6 | | |
| Magnesium acetate | 8 | Cadmium acetate | 7 | | |
| Proline | 5 | Alanine | 5-6 | | |
| Fumarate (di-Sodium salt) | 7 | Threonine | 5 | | |

c

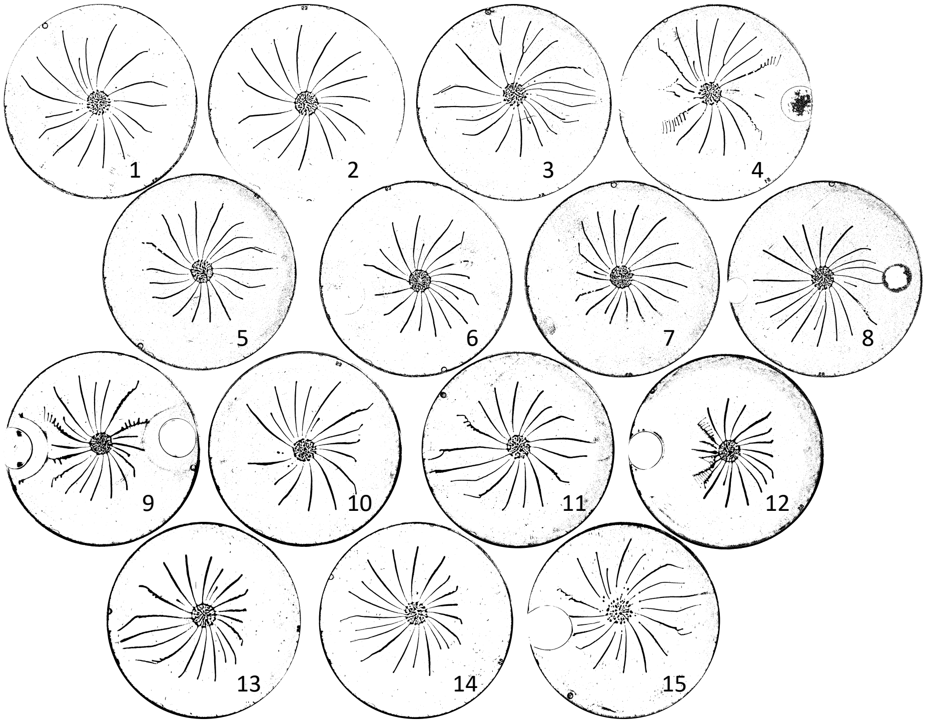
