## Supplementary figures and images for "Cyclic AMP signalling and glucose metabolism mediate pH taxis by African trypanosomes"

### Supplementary Figure 2

#### Supplementary Figure 2

a

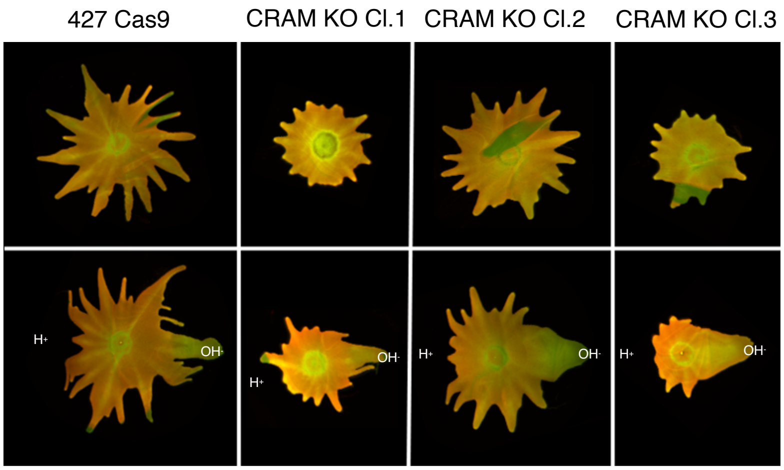


b

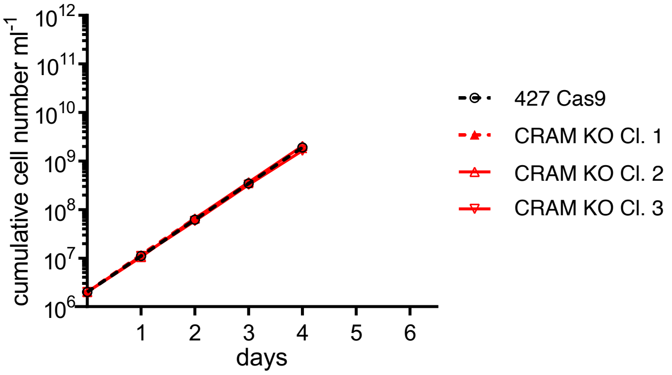

### Supplementary Figure 3

#### Supplementary Figure 3

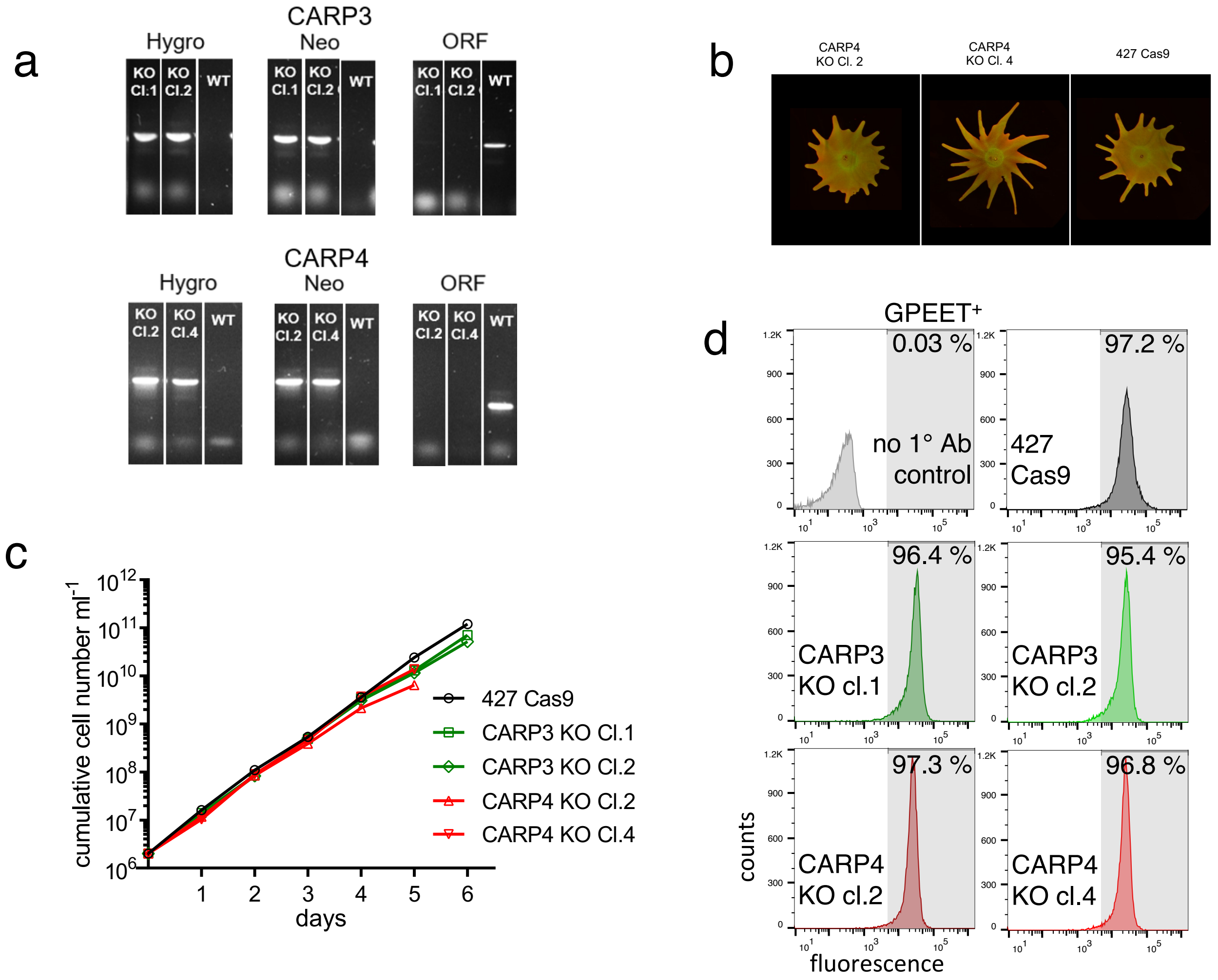

### Supplementary Figure 4

#### Supplementary Figure 4

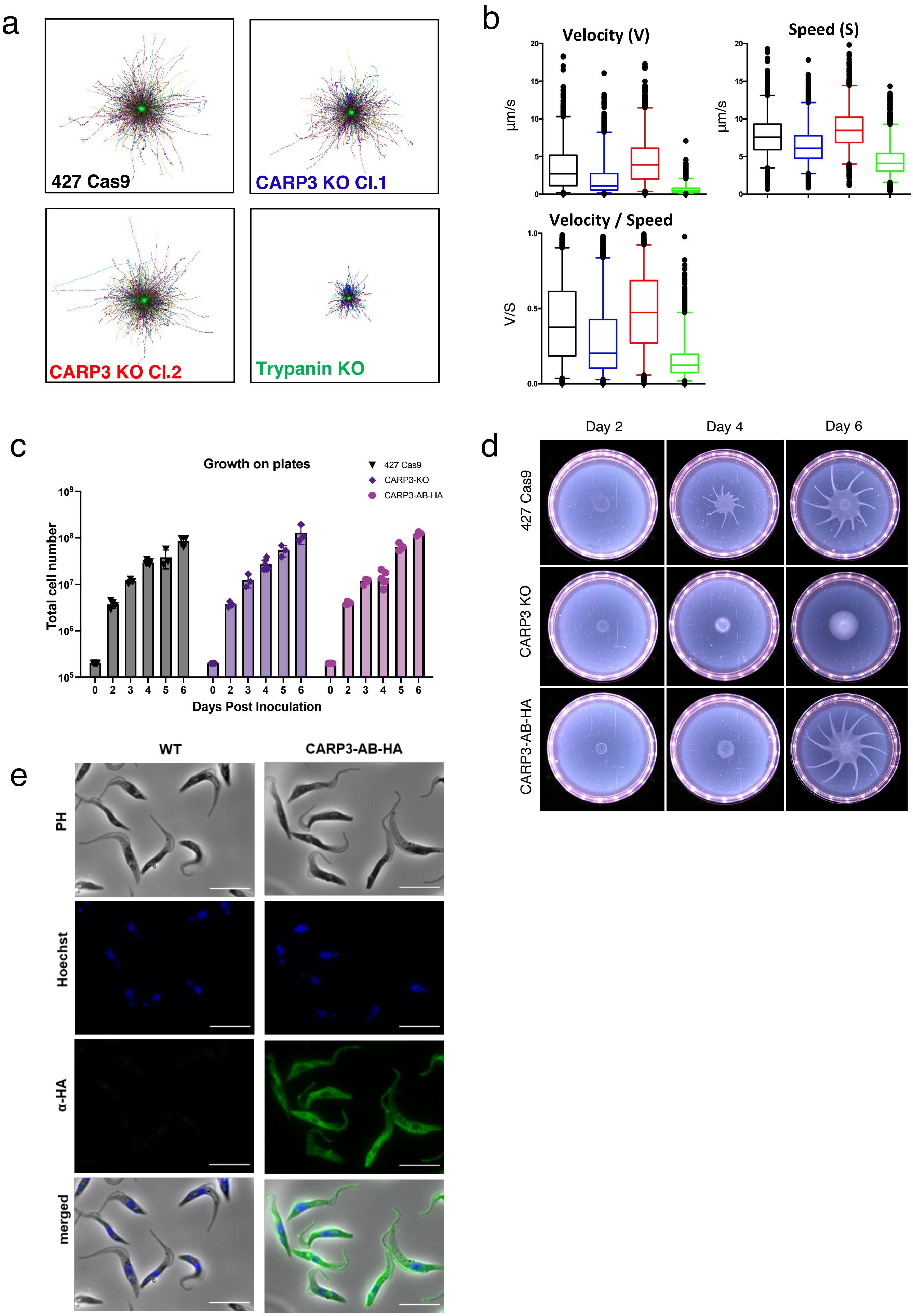

### Supplementary Figure 5

#### Supplementary Figure 5

a

CARP3

Hygro Neo ORF

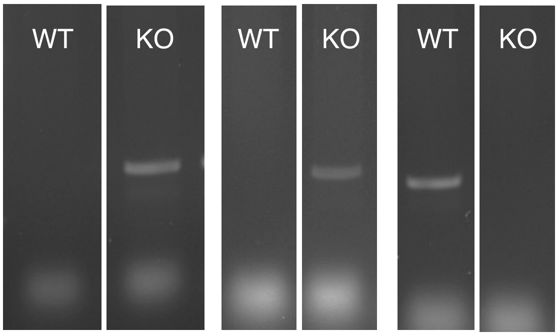


b CARP3 addback

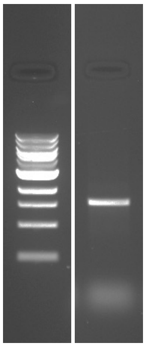

### Supplementary Figure 6

#### Supplementary Figure 6

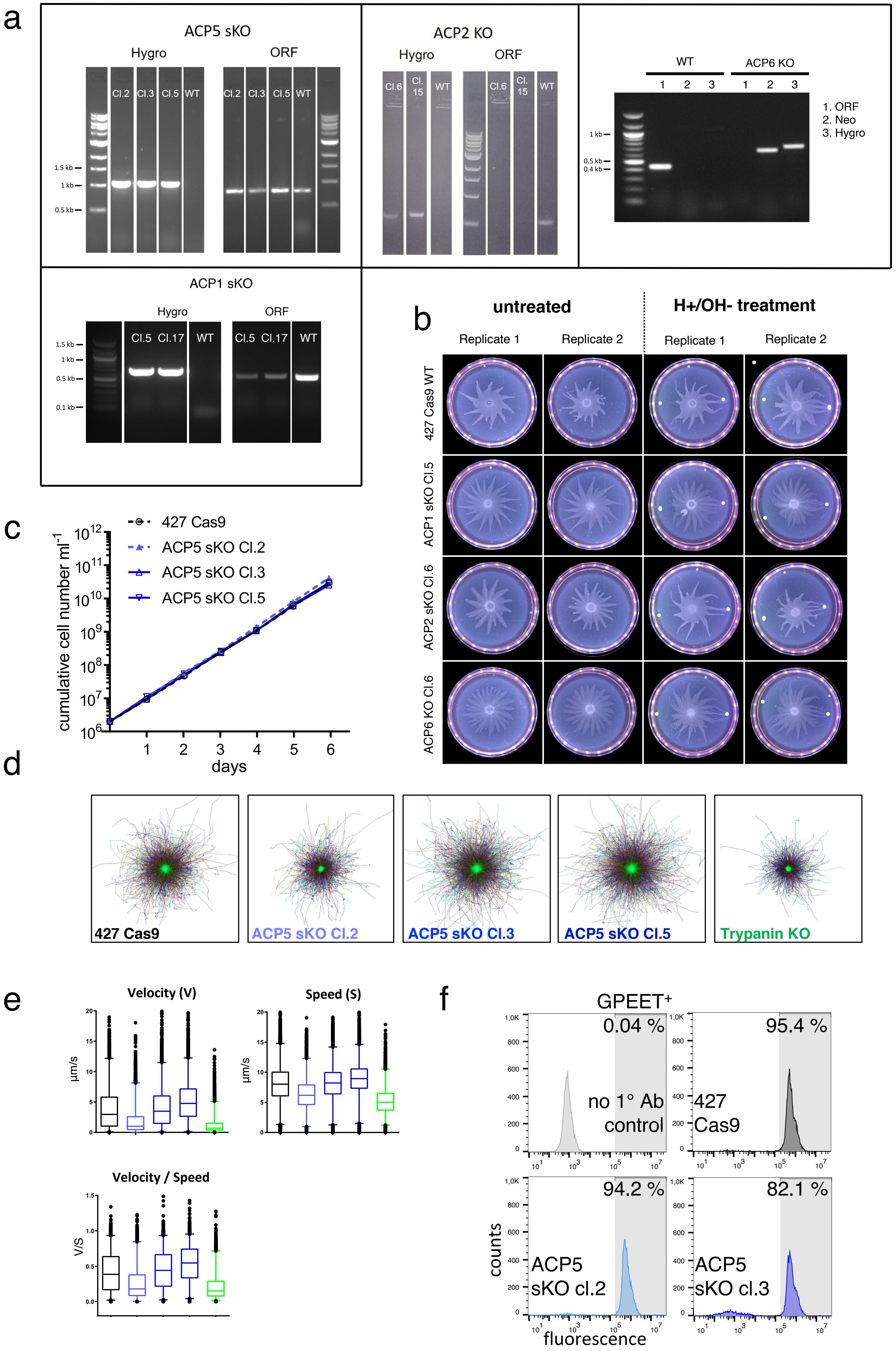
